# Characterization and Heterotic Grouping of Traditional Assam rice (*Oryza sativa* L.)

**DOI:** 10.1101/2021.03.10.434899

**Authors:** Praveen Kumar, Debojit Sarma, Mainu Hazarika

## Abstract

Parents of heterotic hybrids are derived from different heterotic groups with high genetic divergence. Classification of traditional Assam rice germplasm in divergent pools will be advantageous to maximize the heterosis and thereby to ensure food security. In the present investigation, a group of 60 upland rice genotypes were characterized using 53 polymorphic simple sequence repeats (SSR) markers out of 83 molecular markers. The genetic divergence study using unweighted Neighbour-joining (UNJ) method clustered the 60 genotypes into 3 major clusters. The eleven most divergent genotypes identified were crossed in half diallel fashion to determine the mid-parent and better-parent heterosis values for the objective of heterotic grouping. No correlation between heterosis and genetic distance can be attributable to the use of a subset of markers not linked to yield or concerned. In genetic distance based heterotic grouping, the intra-group hybrids were recorded a higher frequency of crosses, grain yield per plant, specific combining ability effect, mid parent heterosis, better parent heterosis and standard parent heterosis value than those of inter-group hybrids. Overall, sn extensive choice of parents with attractive traits constellation leading to increased yield of the hybrids for much better complementation must be stressed along with a substantial hereditary distance for augmentation of yield heterosis.

## Introduction

The global population comprises over 7 billion people, and the global population clock is recording continuous growth (Borem et al. 2014). Rice is the second most important crop in the world, feeding more than one half of the world population (Sharma et al. 2020). Above 90% of rice is consumed by Asians. Today the population growth is the alarming rate, so hybrid rice is a viable and practical approach for feeding tremendously increasing population of India as well as the world. Rice provides almost 15% of worldwide per capita protein requirements (IRRI 2002). The achievement of the hybrid rice that resulted from the exploitation of heterosis is based on the genetic divergence of germplasm, mostly on geographic change for the three-line hybrid rice and sub-specific genetic divergence for the two-line hybrid rice. North-East (NE) India is a traditional home for a vast array of rice germplasm, and high genetic diversity is expected for the landraces belonging to this region (Sahoo et al. 2019). Because of its unique eco-geographical features and broad ethnic diversity, it possesses at least 10,000 indigenous rice landraces that cultivated under lowland, upland, and deep water conditions (Hore et al. 2005).

Rice occupies 4.58 million hectares which are 75% of the total cultivated area of the region (GOI 2015). The rice landraces are grown in diverse ecosystems spreading from the high altitude of Arunachal Pradesh, the food-prone regions of Assam, and rainfed, irrigated, upland, steep terraces and deepwater, Jhum and *tilla* land ecologies. Of the various classes of rice cultivated in Assam, upland rice cultivars of North East India are directly sown in the fields during March-April and harvested in June-July known agronomically as “*aus*/*ahu*” rice (Travis et al. 2015). Genetic diversity among landraces and their genetic relationship is helping in conservation and parental selection in the future breeding programme. Identification of landraces with a high level of genetic variation will be a valuable resource for broadening the genetic base. Since it allows superior alleles to be identified for improved quantitative traits. On the other hand, DNA-based molecular markers are persistent, repeatable, steady, vigorous and profoundly dependable (Virk et al. 2000). Among the accessible DNA markers, simple sequence repeat (SSR) is suitable because of their multi-allelic nature, high reproducibility, co-dominant nature, plenitude and genome-wide coverage (Temnykh et al. 2000), which differ in the level of polymorphism relying upon their area in the coding or non-coding portions, nature of their recurrent themes and the genome-wide coverage. Along these lines, a perfect arrangement of SSR markers giving genome-wide inclusion encourages the measure of the hereditary variation, which provides an intense, unambiguous molecular depiction of rice cultivars/landraces. Previously, numerous scientists have reported the evaluation of the descend varieties of Indian rice germplasm (Sivaranjani et al. 2010; Vanniarajan et al. 2012).

Parents of heterotic hybrids are usually derived from different heterotic groups with high genetic divergence. Grouping of germplasm in divergent pools is advantageous to maximize the expected heterosis (Reif et al. 2005). Heterotic groups for hybrid crops could be determined by marker-based groups as studied in maize (Choukan et al. 2006; Lu et al. 2009; Romay et al. 2013; Xie et al. 2008), chickpea (Sant et al. 1999), rice (Kwon et al. 2002), alfalfa (Riday et al. 2003; Tams et al. 2006) and in maize (Geleta et al. 2004) confirmed the non-significant correlation of genetic distance with heterosis. So, the genetic distance may not be as such reliable for rice hybrid breeding programme and predicting the heterosis at the DNA level. Due to inadequate information on the association of functional molecular markers and yield, the evidence derived from molecular markers at this time is limited to the use of assigning parents into germplasm group or heterotic groups, and to provide a general guideline of avoiding heterotic groups from blending during parent breeding. Full detailed information about functional markers related to yield heterosis may provide precise direction to upsurge the breeding efficiency of rice.

## Material and method

### Experimental materials

A collection of 60 rice genotypes consisting of landraces, maintainers of wild abortive cytoplasmic male sterile (WA-CMS) lines and high yielding varietiesfrom Assam, India were used in the present study. Pedigree and origin of the genotypes are presented in Table 1. The latitude, longitude and altitude of Jorhat are 26°44’N, 94°l0’E and 9l m. above mean sea level respectively. All the molecular work, including DNA extraction, PCR and gel electrophoresis, were performed in the marker laboratory of the Department of Agricultural Biotechnology, Assam Agricultural University, Jorhat. The total genomic DNA from each of the 60 rice genotypes was extracted from 5 g leaves of 21 days old seedlings following the protocol of (Dellaporta et al. 1983) with minor modifications.

### Molecular data analysis

Genetic distance (GD) between each pair of parents was measured (Cavalli-Sforza and Edwards 1967), using PowerMarker version 3.25 (Liu and Muse 2005). Based on Cavalli-Sforza and Edwards chord distances, a dendrogram was constructed illustrating the genetic relationship among the rice genotypes using unweighted Neighbour-joining (UNJ) method as proposed by (Gascuel 1997), which uses a criterion of weighted average, in DARwin 6 (Perrier and Jacquemoud-Collet 2006). The number of alleles per locus, major allele frequency, gene diversity, heterozygosity and polymorphism information content (PIC) values was calculated using PowerMarker version 3.25 (Liu and Muse 2005).

### Cavalli-Sforza chord distance

Cavalli-Sforza and Edwards 1967 proposed the chord distance assuming that the genetic differences arise due to genetic drift only. The chord distance between two populations represented on the surface of a multidimensional hypersphere using allele frequencies at the *j*^*th*^ locus is given by

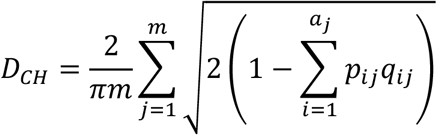

Let *p*_*ij*_ and *q*_*ij*_ be the frequencies of an *i*^*th*^ allele at the *j*^*th*^ locus in populations *X* and *Y*, respectively and*a*_*j*_ is the number of alleles at the *j*^*th*^ locus, and *m* is the number of loci examined? The binary data were used to generate a dissimilarity matrix and genetic diversity using chord distance in PowerMarker version 3.25 (Liu and Muse 2005).

### Gene diversity (He)

The gene diversity of expected heterozygosity (*He*) is defined as the probability that two randomly chosen alleles from the population are different. Gene diversity (Liu and Muse 2005) is defined as follows :

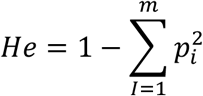

Where, *m* is the number of alleles in a gene locus and *p* is the *i*^*th*^ allele frequency.

### Polymorphic information content (PIC)

The Polymorphic information content (PIC) values provide an estimate of the discriminatory power of the locus or loci by taking into account not only the number of alleles that are expressed but also the relative frequency of those alleles. The informative potential of a marker is high if the PIC value is more than 0.5, moderate if the PIC is between 0.5 and 0.25, and only slightly informative if the PIC value is below 0.25 (Ditta et al. 2018). The polymorphic information content (PIC) for each locus (Botstein et al. 1980) is defined as follows:

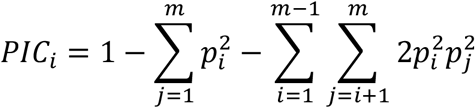

Where, *p*_*i*_ and *p*_*j*_ represent the frequencies of the *i*^*th*^ and the *j*^*th*^ alleles, respectively and m is the number of alleles in a locus.

### Morphological Characters

All the observations except seedling establishment, days to panicle initiation, 50% flowering and maturity were based on 3 random competitive plants. The panicles of the main culms of the sampled plants were cut just below the neck and were separately bagged. Shoot dry weights inclusive of grains for the sample plants were recorded after drying in a hot air oven at 70°C until constant weight. Initial weights of the harvested shoots with intact grains and the separated grains were noted. The filled grains were separated from the remaining shoot portion of each sampled plant before drying to enable moisture determination of the shoots inclusive of grains and the grains alone. The characters namely average panicle weight, straw yield, grain yield and biological yield were reported at 12 per cent moisture content. The observations on flag leaf length/ width were taken at booting stage and on culm length, productive tillers, panicle length was recorded during physiological maturity stage in the standing crop.

#### Seedling height at 21 DAS (SH)

**t**he average length of five random seedlings (shoot portion) was measured in cm at 21 days after sowing (DAS).

#### Leaf number at 21 DAS (LN)

**t**he total number of leaves of five random seedlings were counted at 21 DAS and the average was taken to get leaf number per seedling.

#### The seedling establishment at 7 DAT (SEST)

**t**he seedling establishment was calculated as the ratio of the number of seedlings survived at 7 days after transplanting (DAT) to the total number of seedlings transplanted and expressed in percentage.

#### Days to panicle initiation [DPI]

**d**ays to panicle initiation was recorded from the date of sowing to the date of first panicle emergence in each single row plot as a single observation.

#### Days to 50 per cent flowering [DFF]

**t**he number of days taken from sowing to 50% of the plants in each row with emerged panicles was recorded in days on a plot basis.

#### Flag leaf area (cm^2^) [FLA]

**t**he flag leaf area was calculated at booting stage using the formula given by Yoshida (1981) as follows: Leaf area (cm^2^) = 0.75 × length (cm) × width (cm)

#### Days to maturity [DM]

**i**t was recorded as the number of days taken from sowing to the time when more than 80 per cent of the grains on the panicles of 80 per cent of the plants in a row turned fully yellow.

#### Culm height (cm) [CH]

**t**he height of the main culms of the sampled plants were measured in cm from the ground level to the base of the panicle and the average worked out.

#### The number of productive tillers per plant [PT]

**t**he average number of panicle bearing tillers of the sampled plants per row was recorded at harvest stage.

#### Average panicle weight (g) (APW)

**t**he average weight of the panicles borne on main culms of the sampled plants were recorded in gram before counting the total number of grains.

#### Panicle length [PL]

**t**he average length of the panicles borne on the main culms of five random plants was measured in cm from the panicle neck to the tip of the panicle.

#### The number of filled grains per panicle [GP]

**t**he number of filled grains in the panicles borne on the main culms of the sampled plants was counted and the average worked out.

#### Spikelet fertility (%) [SF]

**t**he spikelet fertility was calculated as the ratio of the number of filled grains to the total number of grains per panicle expressed in percentage.

#### Straw yield per plant (g) [SYP]

**t**he average weight of the shoot portions excluding the grains for the sampled plants was reported as straw yield per plant in gram.

#### Grain yield per plant (g) [GYP]

**t**he average weight of the filled grains of the sampled plants was expressed as grain yield per plant in gram.

#### Biological yield per plant (g) [BYP]

**i**t was calculated as the sum of straw yield and grain yield of each sampled plant and the average expressed in gram.

#### Harvest index (%) [HI]

**t**he harvest index was calculated in percentage as the proportion of grain yield to the biological yield:

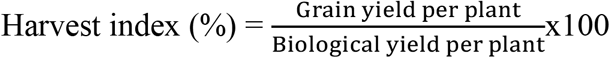

#### Gel consistency [GC]

test was conducted by the method described by (Cogampang et al. 1973). The method separates high amylose rice into hard gel consistency (26-40 mm), medium gel consistency (41-60 mm) and soft gel consistency (61-100 mm) types.

#### Amylose content (%) [AC]

amylose content was determined by using the method described by Dela Cruz and Khush (2000). Rice varieties were grouped based on their amylose content into waxy(0-2%), very low (3-9%), low (10-19%), intermediate (20-25%) and high (>25%)(Kumar and Khush 1986).

#### Heterosis Analysis

The estimates of heterosis were calculated as a percentage increase or decrease of *F*_*1*_s over the mid parent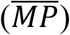, better parent 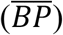 and standard parent 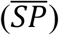 values following the method of (Turner 1953; Hayes et al. 1955).

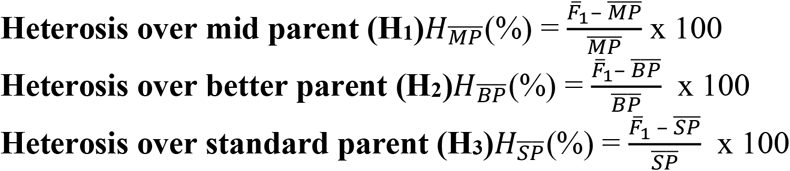

Where, *F*_*1*_ of *F*, 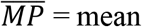 of the two parents, 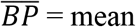 of the better parent and 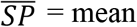 of the standard parent.

**t**he heterosis was tested by the least significant difference at 5% and 1% level of significance for error degree of freedom (*m*) as follows. For testing heterosis over mid parent 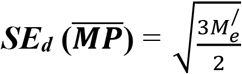

For testing heterosis over better parent and standard parent 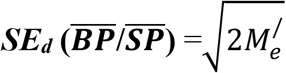

#### Critical difference

CD = SE_d_ x t (at 5 and 1% level for error d.f.), heterosis was considered significant when 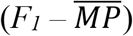 or 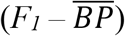 was higher than the critical difference.

## Result and discussion

### Molecular marker analysis

#### Polymorphism of SSR markers

Among the classes of repetitive DNA sequences that have shown acquiescent for PCR amplification, SSRs remains the perfect choice of markers (Jacob et al. 1991). SSR markers are valuable genetic markers because they detect a high level of allelic diversity, and are co-dominant, accessible and economically assayed by PCR (Weber and May 1989). They are easily automated (Westman and Kresovich 1997), abundance and have even genomic distribution and high level of polymorphism (Verma et al. 2019). It has a common polymorphism of at least 1.5 times higher than AFLP and RAPD markers (Mackill et al. 1996). SSRs are highly polymorphic, even between closely related lines (Gupta et al. 1999). The polymorphism in SSR could be due to changes in the SSR region itself caused by the expansion or contraction of SSR or interruption (Li et al. 2000). In the present investigation, a total of 83 SSR markers distributed throughout the 12 chromosomes were used to assess the extent of molecular diversity across 60 rice genotypes of Assam maintained in the Department of Plant Breeding and Genetics of Assam Agricultural University, Jorhat. Out of 83 SSR markers tested, 53 were determined polymorphic.

#### Summary statistics about the SSR markers

DNA bands were scored for DNA fingerprinting analysis with the molecular data generated using 53 SSR markers. Some gel pictures of the polymorphic markers (RM 293, RM 447, RM 429, RM 337, RM 245, RM 152 and RM 216) showed (Supplementary Figure 1). The major allele frequency, number of alleles per locus, gene diversity, heterozygosity and PIC values (Table 2) were calculated for each SSR marker using Power Marker v 3.25 (Liu and Muse 2005). All the genotypes were scored for the allelic weight of the SSR bands throughout all the 60 genotypes. DARwin 6 used an unweighted neighbour method to construct the dendrogram showing the distance-based interrelationship among the genotypes. For the phylogenetic tree, genetic distance was calculated as the chord distance in PowerMarker v 3.25 (Liu and Muse 2005).

#### Major allele frequency

The major allelic frequency revealed by the SSR markers across the 60 rice genotypes ranged from 0.400 to 0.983, with a mean of 0.701. Generally, the allele frequency of a maximum number of markers was below 0.95, indicating that they were all polymorphic. Choudhury et al. 2014, reported major allele frequency ranging from 0.50 for Assam to 0.99 for Tripura. Gour et al. 2017, reported the average of major allele frequency from 0.42 to 0.10 and Mvuyekure et al. 2018, reported 0.76; all these findings were quite similar to our results.

#### Number of alleles

The SSR markers have significantly superior allelic diversity of microsatellites (McCouch et al. 1997) and high numbers of alleles for rice microsatellite markers. In the present study, a total of 133 alleles were detected across 60 rice genotypes by 53 polymorphic SSR markers with an average of 2.5 alleles per locus. The number of alleles generated per locus by each marker ranged from 2 (RM296, RM1896, RM204, RM249, RM12, RM519, RM11, RM20, RM44, RM318, RM530, RM3614, RM5361, RM21, RM202, RM287, RM50, RM520, RM29, RM60, RM261, RM-325, RM28519, RM127, RM171, RM178, RM216, RM222, RM252, RM3331, RM1352, RM15669 and RM438 to 5 (RM260). These numbers were quite comparable to 2.0-5.5 alleles per SSR locus (Cho et al. 2000) using a different set of rice germplasm. A range of 2-4 alleles per locus was reported (Gour et al. 2017; Ming et al. 2015; Mvuyekure et al. 2018; Verma et al. 2019). The average number of alleles was 6.72 (Ravi et al. 2003), 4.90 (Zeng 2004), 4.50 (Pervaiz et al. 2010), 8.00 (Zhao et al. 2009), 4.30 (Islam et al. 2011), 7.40 (Shamim et al. 2015) and 6.40 (Watanabe et al. 2016), which might be due to the use of a more diverse set of rice accessions in their study or due to the use of highly polymorphic markers.

#### Gene diversity

Gene diversity ranged from 0.033 in RM204 to 0.706 in RM334, with a mean of 0.387. Gour et al. 2017 and Mvuyekure et al. 2018, reported mean gene diversity value of 0.390 and 0.325, respectively, which were similar to present mean value. Watanabe et al. 2016, reported a gene diversity mean of 0.84, and Choudhury et al. 2014, obtained gene diversity ranging from 0.006 in Arunachal Pradesh to 0.50 in Manipur.

#### Observed heterozygosity

The observed heterozygosity ranged from 0.017 for RM287 to 0.983 for RM334, with a mean of 0.152. The majority of the SSR markers exhibited observed heterozygosity either zero or low value. Watanabe et al. 2016 and Gour et al. 2017, reported heterozygosity of 0.690 and 0.386, respectively. Choudhury et al. 2014, reported heterozygosity ranging from 0.002 in Nagaland to 0.420 in Mizoram. The present results suggested that the majority of rice germplasm were pure and completely homozygous for SSR markers which might be the result of the self-pollinated mode of reproduction of rice. The observed heterozygosity (0.152) was far lower than total expected heterozygosity (0.387) which further supported a low gene flow value for the majority of the loci, except RM 440, RM 19, RM 260, RM 206, RM 178, RM 293, RM 447 and RM 334.

#### Polymorphism information content (PIC)

In the present investigation, the mean PIC value for all the markers RM 293, RM 334 and RM 545 showed higher discriminatory power to distinguish genotypes due to its high PIC values of 0.645, 0.561 and 0.549, respectively. The primer RM 60 and RM 573 showed lower PIC values 0.090 and 0.064, respectively, suggesting less discriminatory power of these primers. A microsatellite marker having a PIC value of greater than 0.5 is highly informative (Botstein et al. 1980). The average PIC values of 0.33 (Singh et al. 2004), 0.32 (Ming et al. 2015), 0.32 (Gour et al. 2017) and 0.26 (Mvuyekure et al. 2018), reported previously. Pachauri et al. 2013, obtained the mean PIC value of 0.37 in sets of 14 improved varieties and 27 landraces of rice collected from different zones of seven Indian states, which were comparable to the present result. All these findings indicated that the rice accessions used in the present study had a broad genetic diversity. The average PIC values reported were 0.57 (Ravi et al. 2003), 0.57 (Zeng 2004), 0.66 (Zhao et al. 2009), 0.46 (Seetharam et al. 2009), 0.53 (Pervaiz et al. 2010), 0.603 (Islam et al. 2011), 0.57 (Shamim et al. 2015), 0.665 (Yadav et al. 2015), and 0.47 (Anh et al. 2018), respectively. These values were comparatively higher than the earlier reports, which might be due to the high diversity of rice accessions used in their studies or the use of highly polymorphic markers.

The dendrogram showing hierarchical horizontal clustering of 60 *Ahu* rice genotypes of Assam using unweighted neighbour-joining (UNJ) method based on chord distances estimated from 53 polymorphic SSR marker data is presented in Fig. 1. The dendrogram indicated dissimilarity values ranging from 0.111 to 0.695 with 3 major clusters. Cluster I had two sub-clusters I-A and I-B containing 18 and 3 genotypes, respectively. Similarly, cluster II was the largest cluster with 34 genotypes which consisted of two sub-clusters II-A having 16 genotypes and II-B with 18 genotypes. Cluster III contained 5 genotypes belonging to landraces only. All the genotypes of cluster I belonged to landraces except Nagina 22 which was a selection from Rajbhog. Cluster II-A was an admixture composed of both improved varieties and landraces. All the maintainer lines of WA CMS belonged to cluster II A. Bau Murali, and Krishna were the most distantly apart genotypes among all the 60 genotypes. The cluster composition is shown in Table 3.

**Figure 1.**
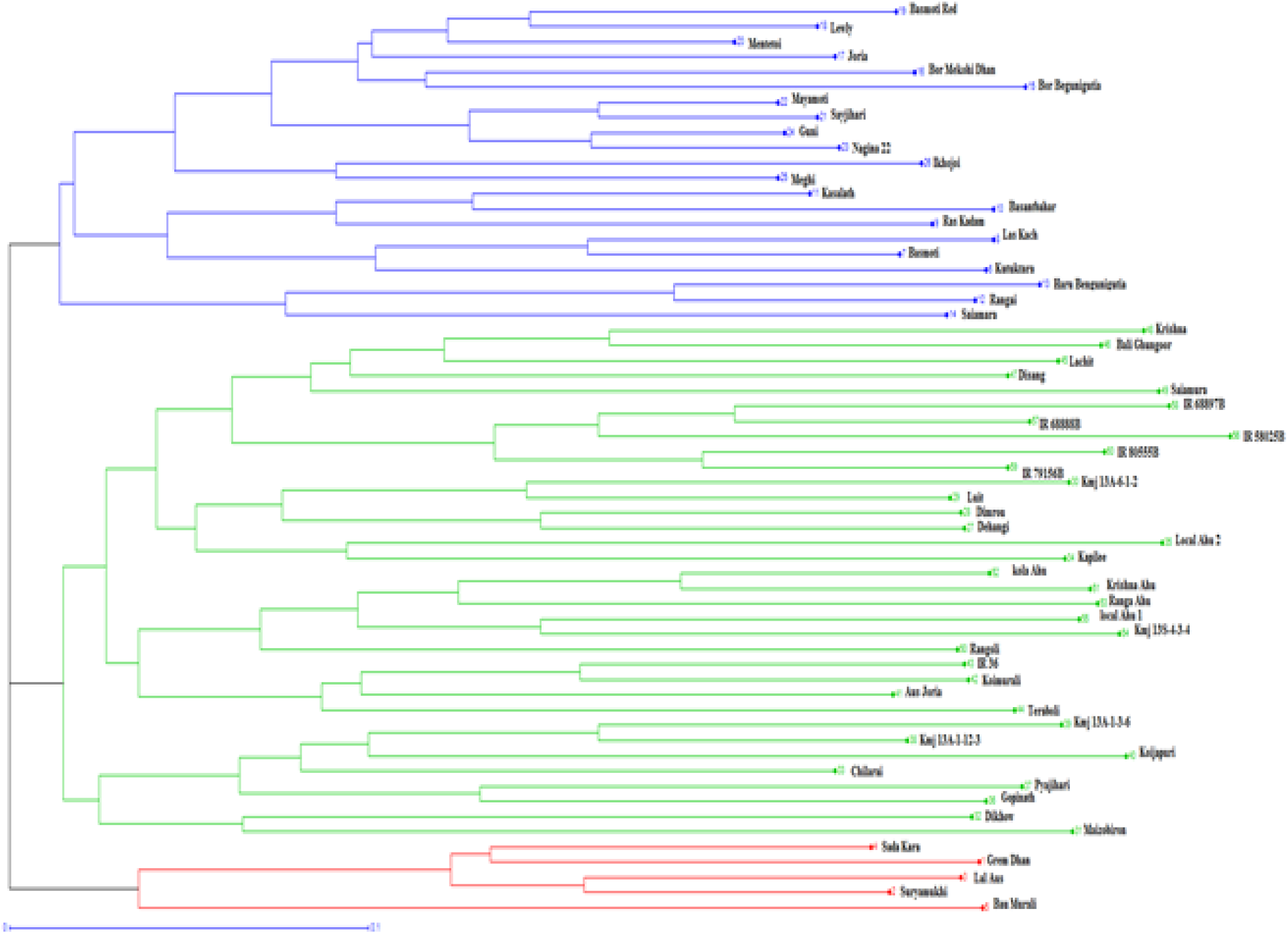
Hierarchical horizontal clustering of 60 *Ahu rice* genotypes of Assam using Unweighted Neighbour-joining (UNJ) method based on chord distances estimated from SSR marker data.

**Fig 1:**
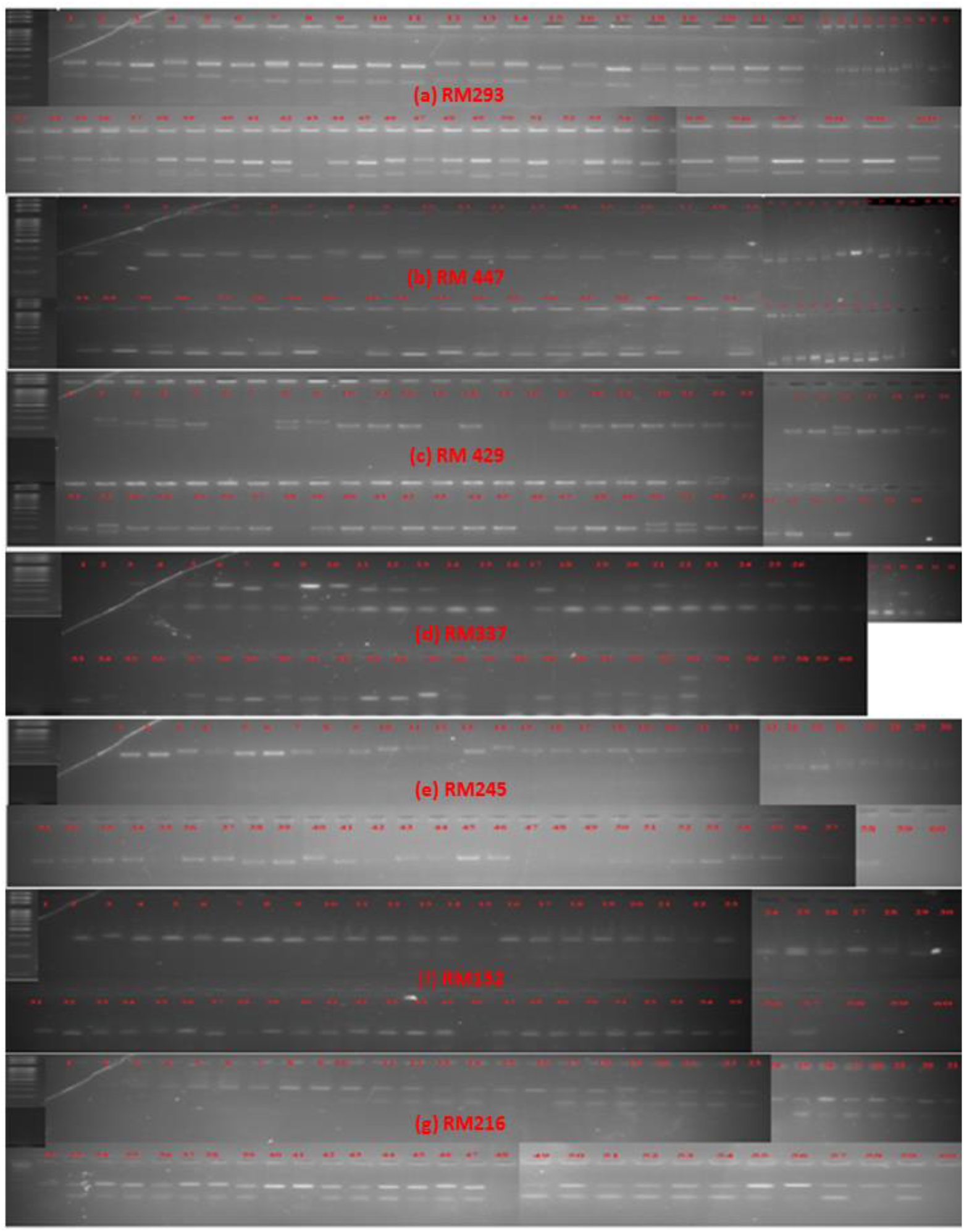
Gel pictures of some of the polymorphic markers showing the amplified products.

Fig. 2. showed 53 SSR markers used for hierarchical clustering using the Unweighted Neighbour-joining (UNJ) method of 11 *Ahu* rice genotypes used in the present investigation. The grouping resulted by using the chord genetic distances among the 11 parental genotypes. All the genotypes included in each cluster represent a heterotic group. As shown in Table 4, three broad heterotic groups were identified using 53 polymorphic SSR markers. Group, I contained BorMekohiDhan and Mayamoti, group II comprised of Luit, Lachit, IR 58025B, IR 68888B, IR 68897B, IR 79156B and IR 80555B, group III included Suryamukhi and Lal Aus. All the maintainers of WA CMS belonged to group II along with Luit and Lachit. Landraces belonged to two different groups I and III. Thus, a fair degree of similarity was evident among genotypes of the same group. Also, the genotypes having a similar combining ability performance and the heterotic response would categorize them into one heterotic group (Melchinger and Gumber 1998).

**Figure 2:**
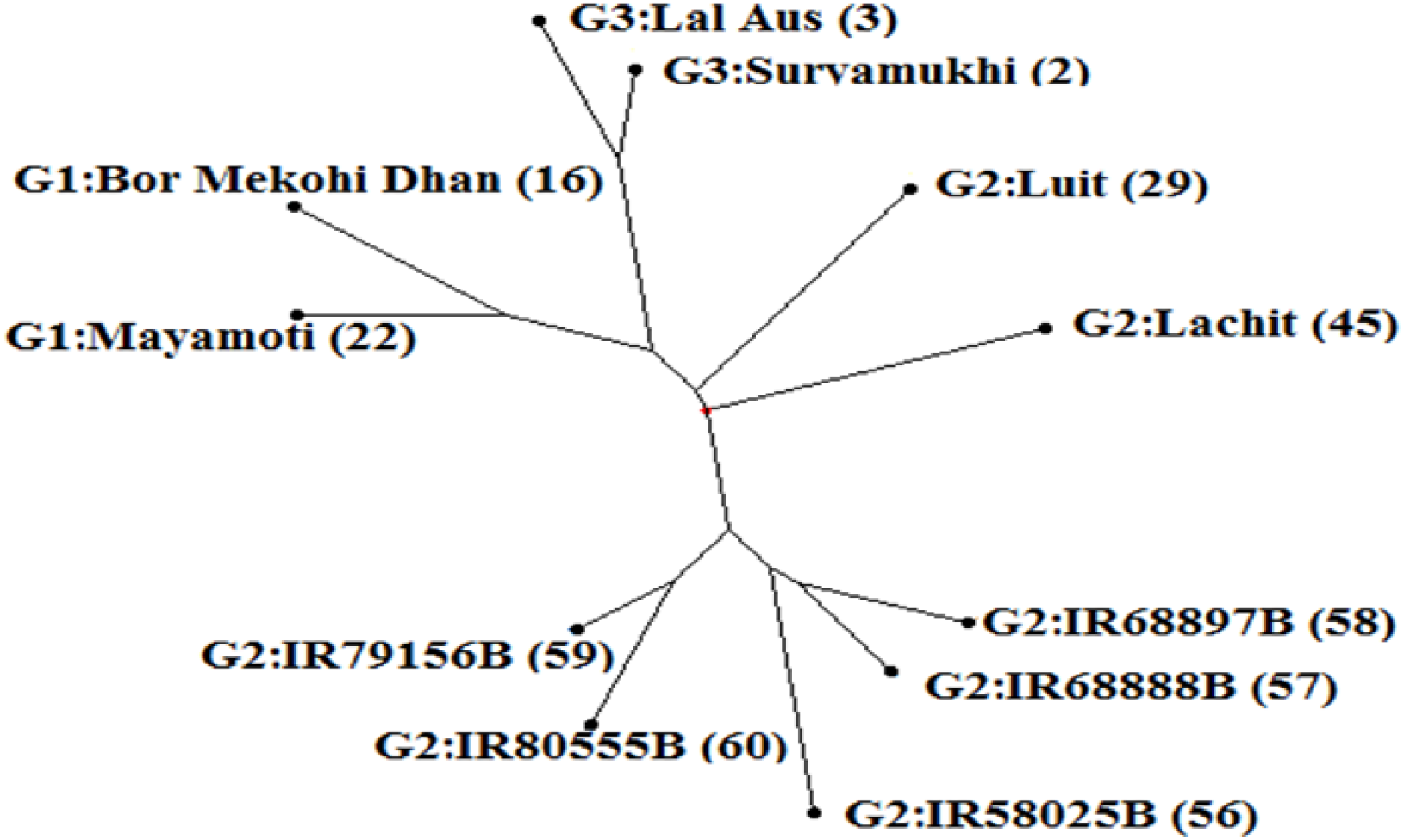
Hierarchical axial clustering of 11 *Ahu rice* genotypes using Unweighted Neighbour-joining (UNJ) method based on genetic distances estimated from SSR markers.

#### Clustering based on genetic distances

The heterotic pattern increases the efficiency of hybrid development, inbred recycling, and population improvement. Recognition and determination of heterotic groups and patterns are essential for breeding hybrid varieties, as shown in various studies on rice (He et al. 2012; Xie et al. 2014). Inbred varieties are then often developed from crosses within heterotic groups. Promising hybrids result from crossing the inbred lines developed between different heterotic groups. Grouping of germplasm in divergent pools is advantageous to maximize the expected heterosis (Reif et al. 2005). The success of the hybrid rice that resulted from the utilization of heterosis depends on the genetic divergence of germplasm, basically on geographic divergence for three-line hybrid rice and sub-specific genetic divergence for two-line hybrid rice. Heterotic groups for hybrid crops could be determined by marker-based groups as studied in maize (Choukan et al. 2006; Romay et al. 2013; Xie et al. 2008), as well as in other crops (Tams et al. 2006). He et al. 2012, investigated the genetic diversity of hybrid rice parents developed at IRRI and evaluated with simple sequence repeats and single-nucleotide polymorphism markers where they confirmed that heterotic hybrids could be formed based on marker-based parent groups to increase the efficiency of hybrid rice breeding (Xie et al. 2014). Fig. 3. showed the frequency distribution of genetic distances (GDs) of the 60 rice genotypes and the selected 11 parents for the diallel crosses based on 53 SSR markers. The mean GD for the 60 genotypes was 0.507 and for the 11 parents was 0.475. This 11 parents’ sample accounted for 83.3% of the allelic variation of the 60 genotypes and reasonably represents the population panel because of the high allelic coverage and similar cluster structure. This result was closely related to Wang et al. 2015.

**Figure 3:**
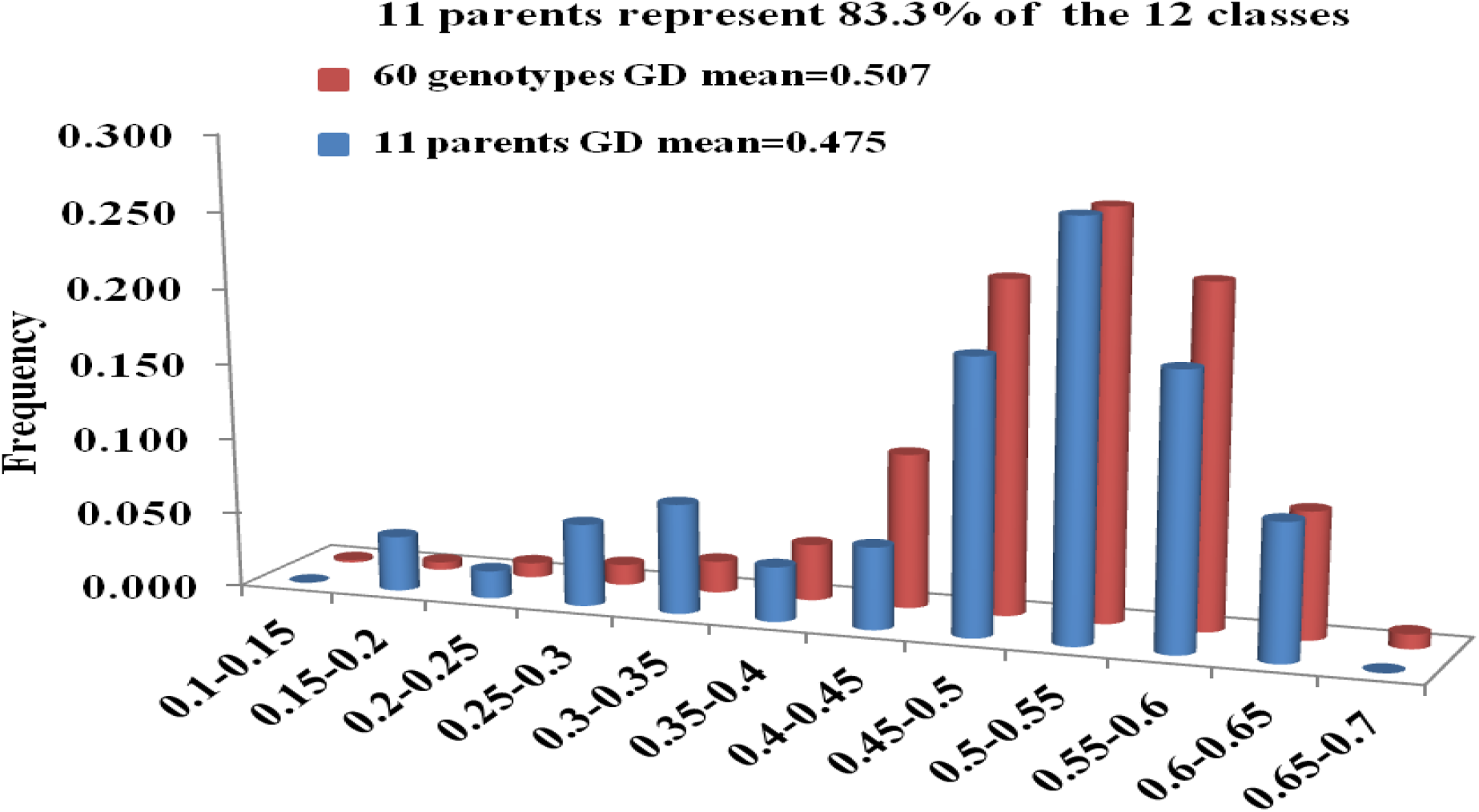
Frequency distribution of genetic distances of the 60 rice genotypes and the selected 11 parents for the diallel crosses based on 53 SSR markers.

#### Pooled analysis of variance (ANOVA)

The pooled analyses of variance (ANOVA) for the various traits evaluated under 40 and 60 kg N ha-1 are presented in **Table 5.1** to **5.3**. The results revealed highly significant (p<0.01) differences among the genotypes for all the traits, indicating that the material under investigation was diverse for the traits in question. The environment source of variation due to Nitrogen (N) doses showed significant differences (p<0.01) for all the traits except only **s**eedling establishment, culm length, panicle length, harvest index, amylose content. The genotype x environment (GE) interaction exhibited highly significant differences (p<0.01) for all traits except **s**eedling establishment, suggesting differential behaviour of the genotypes in the two N doses.

#### Mean comparison

Mean values of the environments (N-doses) averaged over the genotypes for the different traits showing significant environments source of variation are given in Table 5.4 and 5.5. The traits namely, days to panicle initiation (77.7), days to 50% flowering (83.6), flag leaf area (27.1 cm2) days to maturity (111.1), productive tillers plant-1 (16.3), average panicle weight (3.3 g), grains panicle^-1^ (116.1), spikelet fertility (78.0%), straw yield plant^-1^ (62.7 g), grain yield plant-1 (28.5 g), biological yield plant-1 (91.0 g), gel consistency (49.8 mm) obtained for 60 kg N ha^-1^ were significantly higher (P<0.05) than the respective values of the traits at 40 kg N ha^-1^. Thus, increasing the N-dose from 40 to 60 kg had a positive correlation with grain yield and its components traits. Increasing grain yield and biomass could be due to N-supply increase chlorophyll content, leaf area index, and nutrient uptake and utilization. Significant effect of increasing nitrogen rate on rice grain yield was also reported by (Bhuiya et al. 1989; Hussain et al. 1989; Islam et al. 1990; Ebaid and Ghanem 2000; El-Batal et al. 2004; Salem 2006; Wang et al. 2017; Adhikari et al. 2018). Adhikari et al. 2018; Liu et al. 2019, reported that nitrogen application increases the straw yields of transplanted rice. Wang et al. 2017, obtained that N fertilizer doses significantly increased the number of tillers in rice; late-emerging tillers usually produce lower yields compared with early emerging tillers. Mean performances of the parents and the hybrid groups classified as maintainers (M), improved varieties (IV) and landraces (LR) for the traits showing significant genotypic variation are given in **Table 5.6 and 5.7**. Among the two-hybrid groups involving WA CMS maintainers and the remaining parents, M×LR was the most productive for grain yield and other related traits, while LR×IV yielded the maximum with a concomitant increase in component traits. The hybrid group IV×IV was the least productive in terms of yield and most yield component traits, followed by LR×LR and M×M, suggesting that improved varieties were already selected for most desirable alleles for high yield and thus, hybridization among them resulted in less yield improvement, due to few allelic differences between them than crosses between landraces and improved varieties. Given the decrease in genetic diversity among the improved rice varieties due to shifting of the breeding goal from high yield to biotic and abiotic stresses, Singh et al. 2016, suggested broadening the genetic base by incorporating more diverse donor parents in the breeding programme for yield improvement in rice.

#### Range of mid-parent (H_MP_) and better-parent heterosis (H_BP_)

The range of mid-parent heterosis for grains yield per plant was from −6.82** (LAL ×BOR) to 31.11** (88B×LUI) (Table 6). The crosses, BOR×25B, LAL×55B, SUR×LAC and MAY×LAC showed significant positive mid-parent heterosis for grains yield per plant. Significant positive heterosis for grain yield per plant has been reported (Ghara et al. 2012; Kumar and Adilakshmi 2016; Latha et al. 2013; Thorat et al. 2017). Better parent heterosis for grain yield per plant ranged from −10.38** (LAL×BOR) to 29.20** (88B×LUI). The crosses, namely 88B×LUI, SUR×LAC, 88B×55B and MAY×LAC, exhibited significant positive better-parent heterosis for grains yield per plant. A similar finding was also reported by (Sahu et al. 2017).

#### Heterotic grouping based on genetic distances

Genetic distance based heterotic clustering of the 11 parental genotypes along with mean yield, combining ability and heterosis estimates (Table 7) revealed that the intra-group hybrid category G2×G2 contained the highest frequency of crosses (0.38) followed by inter-group hybrid category G1×G2 (0.25) and G2×G3 (0.25). G1×G2 (0.55) registered the highest genetic distance, followed by G1×G3 (0.48) and G2×G2 (0.40). G2×G3 (34.61) recorded the highest grain yield, followed by G3×G3 (33.76) G1×G1 (28.48). G3×G3 (7.03) showed the highest SUM-GCA (g plant-1) followed by G2×G3 (3.05) and G1×G3 (1.63). G1×G1 (5.02) showed the highest SCA (g plant-1) followed by G2×G3 (4.35) and G1×G2 (1.72).

The highest H_MP_ (%) was observed for G2×G3 (63.70) followed by G1×G1 (51.74) and G 3×G3 (46.05). G2×G3 (46.15) exhibited the highest H_BP_ (%), followed by G1×G1 (43.56) and G3×G3 (36.00). G2×G3 (75.60) showed the highest H_SP_ (%), followed by G3×G3 (71.26) and G1×G1 (44.51). G2×G3 had the highest heterosis, grain yield and genetic distance; therefore, hybrids from this group are useful to produce superior hybrids, predicting the heterosis and formed a heterotic pattern. But the highest SCA was shown by G1×G1, an intra-group hybrid category. The lowest yielding hybrids and yield heterosis were evident in the crossing pattern of G1×G3.

The parental group G2 recorded the highest frequency of crosses (0.88), genetic distance (0.50), SCA (2.10), HMP (46.51%) and HBP (32.68%). G3 registered the highest grain yield (31.13), SUM-GCA (3.90) and HSP (57.93%).Mean of the inter-group hybrid categories for the frequency of crosses (0.42), genetic distance (0.53), grain yield (28.74), SUM-GCA (0.78), SCA (0.74), HMP (40.77%), HBP (28.20%) and HSP (45.81%) were 0.42, 0.53, 28.74, 0.78, 0.74, 40.77%, 28.20% and 45.81%, respectively. Mean of the intra-group hybrid categories were 0.58, 0.29, 29.59, 0.78, 1.59, 45.07%, 34.36% and 50.12% for the frequency of crosses, genetic distance, and grain yield, SUM-GCA, SCA, H_MP_, H_BP_ and H_SP_, respectively.

The inter-group parents were more diverse (GD=0.53) than the intra-group parents (GD=0.29), which was in tune with the findings of (Xie et al. 2014; Yingheng et al. 2018). However, the hybrids originating from inter-group parents do not always give high yield or yield heterosis (Wang et al. 2015). But some specific intra-group hybrids, G3×G3 in the present case, having low genetic distance produce a higher yield than the average yield of inter-group hybrids. The present study suggested that molecular markers might be useful for the grouping of parents and heterotic grouping. Based on this, an intra-heterotic group needs genetic improvement, and hybrids between inter-heterotic groups used to produce the superior hybrids. In an intra-group hybrid were observed high grain yield, SCA and heterosis than inter-group hybrids. The hybrid combination of an inter-group giving low yields but have high genetic distance, so genetic distance cannot as such predict the heterosis. However, the use of functional markers related to grain yield and its components would provide specific guidelines for combining parents correctly to increase breeding efficiency. Broadening the hybrid rice parental gene pools by introducing germplasm from other sources and integrating them into heterotic groups are essential steps to further enhance the heterotic performance of hybrids in the state of Assam.

#### Mean comparison of heterotic groups based on genetic distances

Table (8) presents the mean performance of the hybrids in GD-based heterotic groups for the different traits. A comparative evaluation of heterotic group means for different traits could be suggested for improving specific characters. This comparison indicated that G2×G3 had better cluster means for most of the characters and therefore, G2×G3 needs consideration for selecting genotypes as parents in a hybridization programme. The hybrids of G1×G3 group could lead to early maturing hybrid development.

Seedling length (cm), Leaf number, Seedling establishment (%), days to panicle initiation, days to 50% flowering, days to maturity, culm length (cm), productive tillers, grains Panicle-1, harvest index (%) and gel consistency (mm) recorded higher mean performances in the inter-group hybrids than in intra-group hybrids, providing more chances for segregation and recombination and thus, these traits concern priority in selecting promising parents for hybridization programme. For the remaining traits, mean performances of the intra-group hybrids were higher than that of the inter-group hybrids. The higher mean performance was evident for grain yield (g Plant^-1^) in the intra-group hybrids than in the inter-group hybrids and was because of the adverse indirect effects of yield contributing traits on yield in the inter-group hybrids.

#### Correlation between heterosis and genetic distances

A perusal of Table (9) revealed that the parameters, namely, mean yield, HMP, HBP and HSP, were strongly correlated among themselves. Genetic distances showed no correlation with any of the mean yield and heterosis parameters. Grain yield per plant had positive and highly significant correlation (p<0.01) with HMP (g Plant-1), HMP (%), HBP (g Plant^-1^), HBP (%) and HSP(%). HMP (g Plant-1) had positive and significant association with HMP (%),HBP (g Plant^-1^), HBP (%) and HSP (%). HMP (%) showed a positive and significant association with the HBP (g Plant^-1^), HBP (%) and HSP (%). HBP (g Plant^-1^) exhibited positive and significant association with HBP (%) and HSP (%). HBP (%) had a positive and significant association with the HSP (%). The correlation coefficient of genetic distance was negative and non-significant with grain yield per plant (−0.0067). Wang et al. 2015, also found the negative correlation of grain yield with the genetic distance. Singh 2015, found a weak association of grain yield with the genetic distance in maize. Many of the studies on rice found a negative correlation of genetic distance with the heterosis (Singh et al. 2011; Xu et al. 2002; Yu et al. 2011). The association of marker-based GD and hybrid performance was too small to be used for predicting hybrid breeding (Sant et al. 1999; Zhao et al. 2010). Tams et al. 2006 and Geleta et al. 2004, also confirmed the non-significant correlation of genetic distance with heterosis in maize. Thus, the molecular marker-based genetic distance may not always be reliable for rice hybrid breeding programme and prediction of heterosis. The information about functional markers related to yield heterosis might provide precise direction or guideline for combining parents definitely to increase breeding efficiency of rice. Zhang et al. 2010, suggested the prediction of heterosis based on yield-related functional genes. As the heterosis is measured mainly in terms of yield, the reliability of markers within genes (EST-SSRs) in heterosis prediction (Pavani et al. 2018). However, the recent results showed that the association and prediction could be enhanced when the parental groups are formed first by molecular markers, which may not predict the best hybrid combination, but it reveals a practical value of assigning existing and new hybrid rice germplasm into heterotic groups and increasing opportunities to develop desirable hybrids from the best heterotic groups, which is consistent with a previous study in maize (Lanza et al. 1997). Pavani et al. 2018, suggested the use of molecular marker heterozygosity, combining ability, high mean performance for different traits, the morphological and molecular marker-based grouping of parental lines to identify heterotic patterns in rice.

## Conclusions

In genetic distance based heterotic grouping, the intra-group hybrids were recorded a higher frequency of crosses, GYP, SCA, HMP, HBP and HSP value than those of inter-group hybrids. No correlation between heterosis and genetic distance can be attributable to the use of a subset of markers not linked to yield or concerned. Gene linked markers or high-density markers (genome-wide markers) or markers within yield genes should be used for genetic distance analysis and heterotic grouping based on genetic distances. Apart from the further study on the genetic aspects, it might be interesting to integrate epigenomics, metabolomics, proteomics, and systems biology approaches for gaining better understandings into the heterotic gene pools of rice. A careful selection of parents with desirable traits constellation contributing to high yield of the parents for better complementation should be emphasized along with a considerable genetic distance for augmentation of yield heterosis.

## Acknowledgements

We are most thankful to the Department of Agricultural Biotechnology AAU-Jorhat for the SSR genotyping. We also acknowledge the contributions of Dr Debojit Sarma for their helpful comments and writing in the manuscript. Authors are very thankful to anonymous reviewers for their comments and suggestions.

