## Supplementary Tables for "Characterization and Heterotic Grouping of Traditional Assam rice (*Oryza sativa* L.)"

**Table 1: List of the *ahu* rice genotypes of Assam used in the investigation**

| S No. | Name | Origi n | Pedigree | S N | Name | Origi n | Pedigree |
|---|---|---|---|---|---|---|---|
| 1 | GremDhan | Assa | Landrace | 31 | Maizobir | Assa | Landrace |
| 2 | Suryamukhi | Assa | Landrace | 32 | Dikhow | Assa | Heera/Annada |
| 3 | Lal Aus | Assa | Landrace | 33 | Chilarai | Assa | IR 24/CR 44- |
| 4 | Sada Kara | Assa | Landrace | 34 | Kapilee | Assa | Heera/Annada |
| 5 | Bau Murali | Assa | Landrace | 35 | Local | Assa | Landrace |
| 6 | Kutuktara | Assa | Landrace | 36 | Gopinath | Assa | Pusa 2-21/IR |
| 7 | Basmoti | Assa | Landrace | 37 | Pyajihari | Assa | Landrace |
| 8 | Las Kach | Assa | Landrace | 38 | Kmj | Assa | Mahsuri/Luit |
| 9 | Rash Kadam | Assa | Landrace | 39 | Kmj | Assa | Mahsuri/Luit |
| 10 | Basantbahar | Assa | Landrace | 40 | Koijapur | Assa | Landrace |
| 11 | Kasalath | Assa | Landrace | 41 | AusJoria | Assa | Landrace |
| 12 | Rangai | Assa | Landrace | 42 | Koimura | Assa | Landrace |
| 13 | HaruBegunig utia | Assa m | Landrace | 43 | IR 36 | IRRI | IR 1561-228-1 |
| 14 | Saiamara | Assa | Landrace | 44 | Teraboli | Assa | Landrace |
| 15 | BorBegunigu | Assa | Landrace | 45 | Lachit | Assa | CRM 13- |
| 16 | BorMekohiD | Assa | Landrace | 46 | Bali | Assa | Landrace |
| 17 | Joria | Assa m | Landrace | 47 | Disang | Assa m | Heera/Annada |
| 18 | Lewly | Assa m | Landrace | 48 | Krishna E | Assa m | Pure line selection in |
| 19 | Basmoti Red | Assa | Landrace | 49 | Saiamura | Assa | Landrace |
| 20 | Mentetoi | Assa | Landrace | 50 | Rangoli | Assa | Landrace |
| 21 | Sayjihari | Assa | Landrace | 51 | Krishna | Odis | GEB 24/TN-1 |
| 22 | Mayamoti | Assa | Landrace | 52 | Kola | Assa | Landrace |
| 23 | Nagina 22 | Odis ha | Selection from | 53 | Ranga Ahu | Assa m | Landrace |
| 24 | Guni | Assa | Landrace | 54 | Kmj | Assa | Mahsuri/Malb |
| 25 | Meghi | Assa | Landrace | 55 | Local | Assa | Landrace |
| 26 | Ikhojoi | Assa | Landrace | 56 | IR | IRRI | Maintainer |
| 27 | Dehangi | Assa m | Maibee R/CRM49//Maibee | 57 | IR 68888B | IRRI | Maintainer line |
| 28 | Dimrou | Assa | Landrace | 58 | IR | IRRI | Maintainer |
| 29 | Luit | Assa | Heera/Annada | 59 | IR | IRRI | Maintainer |
| 30 | Kmj 13A-6- | Assa | Mahsuri/Luit | 60 | IR | IRRI | Maintainer |

**Table 2: List of polymorphic markers with their allele frequencies, no of alleles, gene diversity, heterozygosity and PIC values**

| S. No. | Marker | Major allele frequency | No. of alleles | Gene diversity (He) | Heterozygosity | PIC |
|---|---|---|---|---|---|---|
| 1 | RM-204 | 0.754 | 2 | 0.371 | 0.017 | 0.302 |
| 2 | RM-164 | 0.604 | 4 | 0.493 | 0.075 | 0.389 |
| 3 | RM-249 | 0.575 | 2 | 0.489 | 0.017 | 0.369 |
| 4 | RM-440 | 0.517 | 3 | 0.576 | 0.433 | 0.489 |
| 5 | RM-12 | 0.767 | 2 | 0.357 | 0.259 | 0.293 |
| 6 | RM-19 | 0.712 | 3 | 0.431 | 0.508 | 0.368 |
| 7 | RM-235 | 0.558 | 3 | 0.520 | 0.350 | 0.415 |
| 8 | RM-260 | 0.536 | 5 | 0.593 | 0.745 | 0.523 |
| 9 | RM-519 | 0.592 | 2 | 0.483 | 0.017 | 0.366 |
| 10 | RM-11 | 0.764 | 2 | 0.361 | 0.000 | 0.296 |
| 11 | RM-20 | 0.733 | 2 | 0.391 | 0.000 | 0.315 |
| 12 | RM-44 | 0.567 | 2 | 0.491 | 0.033 | 0.371 |
| 13 | RM-318 | 0.983 | 2 | 0.033 | 0.000 | 0.032 |
| 14 | RM-530 | 0.767 | 2 | 0.358 | 0.000 | 0.294 |
| 15 | RM-3614 | 0.533 | 2 | 0.498 | 0.000 | 0.374 |
| 16 | RM-5361 | 0.550 | 2 | 0.495 | 0.000 | 0.372 |
| 17 | RM-206 | 0.500 | 4 | 0.585 | 0.933 | 0.501 |
| 18 | RM-252 | 0.950 | 2 | 0.095 | 0.000 | 0.090 |
| 19 | RM-21 | 0.508 | 2 | 0.500 | 0.117 | 0.375 |
| 20 | RM-167 | 0.900 | 3 | 0.182 | 0.033 | 0.168 |
| 21 | RM-202 | 0.867 | 2 | 0.231 | 0.033 | 0.204 |
| 22 | RM-287 | 0.517 | 2 | 0.499 | 0.033 | 0.375 |
| 23 | RM-50 | 0.897 | 2 | 0.185 | 0.000 | 0.168 |
| 24 | RM-520 | 0.850 | 2 | 0.255 | 0.000 | 0.222 |
| 25 | RM-29 | 0.683 | 2 | 0.433 | 0.033 | 0.339 |
| 26 | RM-60 | 0.950 | 2 | 0.095 | 0.000 | 0.090 |
| 27 | RM-261 | 0.867 | 2 | 0.231 | 0.000 | 0.204 |
| 28 | RM-273 | 0.842 | 3 | 0.271 | 0.083 | 0.242 |
| 29 | RM-325 | 0.858 | 2 | 0.243 | 0.050 | 0.214 |

**Table 2contd…**

| S No. | Marker | Major allele frequency | No. of alleles | Gene diversity | Heterozygosity | PIC |
|---|---|---|---|---|---|---|
| 30 | RM-545 | 0.442 | 3 | 0.627 | 0.017 | 0.549 |
| 31 | RM-573 | 0.967 | 3 | 0.065 | 0.000 | 0.064 |
| 32 | RM-1038 | 0.658 | 3 | 0.508 | 0.083 | 0.455 |
| 33 | RM-3331 | 0.85 | 2 | 0.255 | 0.000 | 0.222 |
| 34 | RM-3742 | 0.783 | 3 | 0.352 | 0.367 | 0.309 |
| 35 | RM-28519 | 0.867 | 2 | 0.231 | 0.000 | 0.204 |
| 36 | RM-127 | 0.575 | 2 | 0.489 | 0.017 | 0.369 |
| 37 | RM-171 | 0.783 | 2 | 0.339 | 0.000 | 0.282 |
| 38 | RM-178 | 0.558 | 2 | 0.493 | 0.883 | 0.372 |
| 39 | RM-216 | 0.725 | 2 | 0.399 | 0.017 | 0.319 |
| 40 | RM-222 | 0.767 | 2 | 0.358 | 0.000 | 0.294 |
| 41 | RM-293 | 0.400 | 4 | 0.700 | 0.983 | 0.645 |
| 42 | RM-438 | 0.983 | 2 | 0.033 | 0.000 | 0.032 |
| 43 | RM-496 | 0.408 | 4 | 0.706 | 0.600 | 0.655 |
| 44 | RM-1352 | 0.917 | 2 | 0.153 | 0.000 | 0.141 |
| 45 | RM-15669 | 0.917 | 2 | 0.153 | 0.000 | 0.141 |
| 46 | RM-152 | 0.483 | 3 | 0.546 | 0.000 | 0.442 |
| 47 | RM-245 | 0.517 | 3 | 0.555 | 0.000 | 0.458 |
| 48 | RM-296 | 0.792 | 2 | 0.330 | 0.017 | 0.275 |
| 49 | RM-1896 | 0.608 | 2 | 0.477 | 0.017 | 0.363 |
| 50 | RM-429 | 0.850 | 3 | 0.266 | 0.150 | 0.250 |
| 51 | RM-447 | 0.475 | 3 | 0.636 | 0.717 | 0.564 |
| 52 | RM-347 | 0.558 | 3 | 0.500 | 0.017 | 0.383 |
| 53 | RM-334 | 0.550 | 4 | 0.614 | 0.400 | 0.561 |
| **Mean** | | **0.701** | **2.5** | **0.387** | **0.152** | **0.323** |

**Table3 : Cluster composition of the 60 rice genotypes based on SSR markers**

| Cluster | No. of genotypes | Composition |
|---|---|---|
| **I A** | 18 | Basmoti Red, Lewly, Mentetoi, Joria, BorMekohiDhan, BorBegunigutia, Mayamoti, Sayjihari, Guni, Nagina 22, Ikhojoi, Meghi, Kasalath, Basantbahar, Rash Kadam, Lal Kach, Rangai, Kutuktara |
| **I B** | 3 | HaruBegunigutia, Rangai, Saiamara |
| **II A** | 16 | Krishna E, Bali Ghungoor, Lachit, Disang, Saiamura, Kmj 13A-6-1-2, Luit, Dimrou, Dehangi, Local Ahu 2, Kapilee, IR 58025B, IR 68888B, IR 68897B, IR 79156B, IR 80555B |
| **II B** | 18 | Kola Ahu, Krishna, Ranga Ahu, Local Ahu 1, Kmj 14S-4-3-4, Rangoli, IR 36, Koimurali, AusJoria, Teraboli, Kmj 13A-1-3-6, Kmj 13A-1-12-3, Koijapuri, Chilarai, Pyajihari, Gopinath, Dikhow, Maizobiron |
| **III** | 5 | Sada Kara, GremDhan, Lal aus, Suryamukhi, Bau Murali |

**Table 4: Composition of the heterotic groups obtained from GD-based cluster analysis**

| Group | GD-based |
|---|---|
| I | BorMekohiDhan (16), Mayamoti (22) |
| 2 | Luit (29), Lachit (45), IR 58025B (56), IR 68888B (57), IR 68897B (58), IR 79156B (59), IR 80555B (60) |
| 3 | Suryamukhi (2), Lal Aus (3) |

**Table5.1: Pooled ANOVA for the traits of the 66 genotypes including 11 parents and their 55 hybrids evaluated over the two nitrogen doses.**

| Source of Variations | DF | Mean Squares | | | | | |
|---|---|---|---|---|---|---|---|
| | | **Seedling length (cm)** | **Leaf number** | **Seedling establishment (%)** | **Days to panicle initiation** | **Days to 50% flowering** | **Flag leaf area ($cm^2$)** |
| Replicates/N Doses | 2 | 0.03 (1) | 0.02 (1) | 9.85 | 0.76 | 0.47 | 6.72 |
| Nitrogen Doses (N | 1 | - | - | 151.52 | 992.97** | 1605.31** | 1052.00** |
| Genotypes (Gen) | 65 | 33.16** | 0.61** | 488.25** | 62.59** | 64.69** | 46.92** |
| N Doses×Gen | 65 | - | - | 95.36 | 18.54** | 16.68** | 16.17** |
| Pooled Error | 13 | 1.91 (65) | 0.20 (65) | 82.16 | 2.94 | 2.54 | 3.07 |
| CV (%) | | 6.63 | 15.32 | 11.36 | 2.27 | 1.96 | 1.61 |

Figures in parentheses are degrees of freedom for replication and error, respectively for the traits recorded in the seedling stage. *, ** Significant at 5% and 1% level

**Table 5.2: Pooled ANOVA for the traits of the 66 genotypes including 11 parents and their 55 hybrids evaluated over the two nitrogen doses.**

| Source of Variations | DF | Mean Squares | | | | | |
|---|---|---|---|---|---|---|---|
| | | **Days to maturity** | **Culm height (cm)** | **Productive tillers $Plant^{-1}$** | **Average panicle weight (g)** | **Panicle length (cm)** | **Filled grains $Panicle^{-1}$** |
| Replicates/N Doses | 2 | 47.31 | 11.12 | 5.17 | 0.21 | 0.43 | 11.95 |
| Nitrogen Doses (N | 1 | 62.06** | 21.31 | 293.80** | 9.58** | 0.21 | 10363.81** |
| Genotypes (Gen) | 65 | 57.46** | 1173.89** | 95.43** | 5.07** | 41.17** | 9046.83** |
| N Doses×Gen | 65 | 20.64** | 133.20** | 30.24** | 1.85** | 21.36** | 709.23** |
| Pooled Error | 13 | 5.62 | 20.39 | 2.46 | 0.26 | 3.35 | 11.09 |
| CV (%) | | 2.20 | 6.54 | 10.29 | 16.45 | 7.21 | 3.03 |

*, ** Significant at 5% and 1% level

**Table: 5.3: Pooled ANOVA for the traits of the 66 genotypes including 11 parents and their 55 hybrids evaluated over the two nitrogen doses.**

| Source of Variations | DF | Mean Squares | | | | | |
|---|---|---|---|---|---|---|---|
| | | **Spikelet fertility (%)** | **Straw yield Plant$^{-1}$ (g)** | **Grain yield plant$^{-1}$ (g)** | **Biological yield Plant$^{-1}$** | **Harvest index (%)** | A[illegible] |
| Replicates/N Doses | 2 | 1.42 | 57.93 | 4.52 | 25.80 | 5.78 | |
| Nitrogen Doses (N Doses) | 1 | 110.80** | 2176.21** | 448.58** | 4444.81** | 0.02 | |
| Genotypes (Gen) | 65 | 778.82** | 1276.49** | 383.76** | 1741.82** | 357.12** | |
| N Doses×Gen | 65 | 96.83** | 56.74** | 46.27** | 124.79** | 39.70** | |
| Pooled Error | 130 | 1.78 | 16.99 | 4.24 | 24.93 | 5.38 | |
| CV (%) | | 1.72 | 6.89 | 7.57 | 5.74 | 7.35 | |

*, ** Significant at 5% and 1% level

**Table 5.4: Mean performance of the two N-doses for the traits showing significant environmental variation.**

| N-Dose (kg $ha^{-1}$) | Days to panicle initiation | Days to 50% flowering | Flag leaf area ($cm^2$) | Days to maturity | Productive tillers $Plant^{-1}$ | Average panicle weight (g) |
|---|---|---|---|---|---|---|
| 40 | $73.8^b$ | $78.7^b$ | $26.7^b$ | $107.1^b$ | $14.2^b$ | $2.9^b$ |
| 60 | $77.7^a$ | $83.6^a$ | $27.1^a$ | $111.1^a$ | $16.3^a$ | $3.3^a$ |
| **CD** | **0.4** | **0.4** | **0.2** | **0.4** | **0.4** | **0.1** |

Mean values with different superscript lowercase letters indicate significant difference at the 0.05 level.

**Table 5.5: Mean performance of the two N-doses for the traits showing significant environmental variation.**

| N-Dose (kg $ha^{-1}$) | Filled grains $Panicle^{-1}$ | Spikelet fertility (%) | Straw yield $Plant^{-1}$ | Grain yield $Plant^{-1}$ | Biological yield $Plant^{-1}$ | Gel consistency (mm) |
|---|---|---|---|---|---|---|
| 40 | $103.5^b$ | $76.7^b$ | $56.9^b$ | $25.9^b$ | $82.8^b$ | $44.9^b$ |
| 60 | $116.1^a$ | $78.0^a$ | $62.7^a$ | $28.5^a$ | $91.0^a$ | $49.8^a$ |
| **CD** | **0.8** | **0.3** | **1.0** | **0.5** | **1.2** | **0.7** |

Mean values with different superscript lowercase letters indicate significant difference at the 0.05 level.

**Table 5.6: Mean performance of the parent and the hybrid groups classified as maintainers (M), improved varieties (IV) and landraces (LR) for the traits showing significant genotypic variation.**

| Parent /Hybrid group | Seedling length (cm) | Leaf number | Seedling establishment (%) | Days to panicle initiation | Days to 50% flowering | Flag leaf area (cm$^{2}$) | Days to maturity | Culm height (cm) | Productive tillers Plant$^{-1}$ | Average panicle weight (g Plant$^{-1}$) |
|---|---|---|---|---|---|---|---|---|---|---|
| M | 16.7$^{f}$ | 4.0$^{a}$ | 82.0$^{bc}$ | 68.4$^{f}$ | 73.4$^{f}$ | 24.1 | 103.4 | 48.1 | 12.1$^{f}$ | 2.7$^{cde}$ |
| IV | 18.4$^{def}$ | 3.0$^{bc}$ | 93.8$^{ab}$ | 72.9$^{e}$ | 77.9e | 28.5 | 106.6 | 57.1 | 13.8$^{def}$ | 2.3$^{de}$ |
| LR | 24.4$^{ab}$ | 2.9$^{bc}$ | 84.4$^{bc}$ | 77.1$^{bc}$ | 81.8$^{bc}$ | 24.6 | 109.9 | 76.4 | 14.3$^{cdef}$ | 2.6$^{cde}$ |
| **Parents** | **20.0$^{cde}$** | **3.5$^{ab}$** | **85.0$^{bc}$** | **72.4$^{e}$** | **77.3e** | **25.1** | **106.3** | **60.0** | **13.2$^{ef}$** | **2.6$^{cde}$** |
| M×M | 17.9$^{ef}$ | 2.9$^{bc}$ | 76.8$^{c}$ | 74.0$^{de}$ | 79.5$^{de}$ | 29.4 | 107.3 | 57.9 | 16.5$^{bc}$ | 3.0$^{bcd}$ |
| M×IV | 21.9$^{bc}$ | 2.5$^{c}$ | 80.3$^{c}$ | 75.5$^{cd}$ | 80.9$^{cd}$ | 25.0 | 108.9 | 54.6 | 12.5$^{f}$ | 3.7$^{ab}$ |
| M×LR | 21.1$^{cd}$ | 3.0$^{bc}$ | 77.9$^{c}$ | 78.0$^{b}$ | 83.5$^{b}$ | 26.7 | 111.3 | 78.9 | 16.5$^{bc}$ | 3.2$^{abc}$ |
| LR×LR | 22.7$^{bc}$ | 2.7$^{bc}$ | 81.3$^{bc}$ | 75.8$^{bcd}$ | 81.2$^{cd}$ | 28.2 | 108.2 | 74.9 | 12.1$^{f}$ | 3.8$^{a}$ |
| IV×IV | 27.0$^{a}$ | 3.0$^{bc}$ | 100.0$^{a}$A$^{a}$ | 80.5$^{a}$ | 87.5$^{a}$ | 25.8 | 116.3 | 48.1 | 16.6$^{b}$ | 2.1$^{e}$ |
| LR×IV | 22.0$^{bc}$ | 2.9$^{bc}$ | 76.9$^{c}$ | 76.3$^{bcd}$ | 82.5$^{bc}$ | 28.2 | 110.0 | 87.1 | 18.9$^{a}$ | 2.5$^{cde}$ |
| **Hybrids** | **21.1$^{cd}$** | **2.8$^{bc}$** | **78.7$^{c}$** | **76.4$^{bcd}$** | **82.0$^{bc}$** | **27.3** | **109.7** | **70.9** | **15.6$^{bcd}$** | **3.2$^{abc}$** |
| P + H | **20.9$^{cd}$** | **2.9$^{bc}$** | **79.8$^{c}$** | **75.7$^{bcd}$** | **81.2$^{cd}$** | **26.9** | **109.1** | **69.1** | **15.2$^{bcd}$** | **3.1$^{abc}$** |
| **CD (5%)** | **2.8** | **0.9** | **12.7** | **2.4** | **2.2** | **1.4** | **2.5** | **6.3** | **2.2** | **0.7** |

Mean values with different superscript lowercase letters indicate significant difference at the 0.05 level.

**Table 5.7: Mean performance of the parent and the hybrid groups classified as maintainers (M), improved varieties (IV) and landraces (LR) for the traits showing significant genotypic variation.**

| Parent /Hybrid group | Panicle length (cm) | Filled grains Panicle$^{-1}$ | Spikelet fertility (%) | Straw yield (g Plant$^{-1}$) | Grain yield (g Plant$^{-1}$) | Biological yield (g Plant$^{-1}$) | Harvest index (%) | Amylose content (%) | Gel consistency (mm) |
|---|---|---|---|---|---|---|---|---|---|
| M | 23.8$^{a}$ | 99.7$^{d}$ | 80.8$^{b}$ | 45.4$^{f}$ | 19.9$^{e}$ | 65.4$^{d}$ | 31.1$^{cd}$ | 14.4$^{abc}$ | 48.0$^{cd}$ |
| IV | 24.7$^{a}$ | 81.9$^{f}$ | 88.0$^{a}$ | 53.6$^{de}$ | 18.6$^{e}$ | 72.2$^{cd}$ | 25.5$^{e}$ | 15.9$^{abc}$ | 63.9$^{a}$ |
| LR | 25.0$^{a}$ | 89.4$^{e}$ | 77.4$^{c}$ | 56.0$^{cde}$ | 20.9$^{e}$ | 77.0$^{c}$ | 27.2$^{e}$ | 13.5$^{c}$ | 63.1$^{a}$ |
| **Parents (P)** | **24.4$^{a}$** | **92.7$^{e}$** | **80.8$^{b}$** | **50.8$^{ef}$** | **20.0$^{e}$** | **70.9$^{cd}$** | **28.7$^{de}$** | **14.4$^{abc}$** | **56.3$^{b}$** |
| M×M | 25.3$^{a}$ | 109.4$^{c}$ | 70.6$^{e}$ | 65.1$^{ab}$ | 28.7$^{bc}$ | 92.4$^{ab}$ | 32.1$^{abc}$ | 16.4$^{a}$ | 43.4$^{e}$ |
| M×IV | 25.9$^{a}$ | 80.8$^{f}$ | 76.4$^{c}$ | 69.8$^{a}$ | 25.1$^{d}$ | 94.9$^{a}$ | 25.9$^{e}$ | 13.7$^{bc}$ | 38.6$^{f}$ |
| M×LR | 26.0$^{a}$ | 113.1$^{c}$ | 76.6$^{c}$ | 57.7$^{cd}$ | 30.1$^{ab}$ | 88.1$^{ab}$ | 34.6$^{ab}$ | 14.7$^{abc}$ | 50.0$^{c}$ |
| LR×LR | 25.6$^{a}$ | 142.5$^{a}$ | 74.3$^{d}$ | 61.2$^{bc}$ | 27.1$^{cd}$ | 88.3$^{ab}$ | 31.0$^{cd}$ | 15.1$^{abc}$ | 42.3$^{ef}$ |
| IV×IV | 23.6$^{a}$ | 112.7$^{c}$ | 75.8$^{cd}$ | 52.9$^{de}$ | 19.2$^{e}$ | 72.1$^{cd}$ | 26.5$^{e}$ | 16.1$^{ab}$ | 42.5$^{e}$ |
| LR×IV | 25.0$^{a}$ | 136.7$^{b}$ | 86.6$^{a}$ | 58.1$^{cd}$ | 31.9$^{a}$ | 90.2$^{ab}$ | 35.2$^{a}$ | 13.9$^{bc}$ | 48.2$^{cd}$ |
| **Hybrids (H)** | **25.6$^{a}$** | **113.2$^{c}$** | **76.7$^{c}$** | **61.6$^{bc}$** | **28.7$^{bc}$** | **90.2$^{ab}$** | **32.1$^{abc}$** | **14.8$^{abc}$** | **45.5$^{de}$** |
| P + H | **25.4$^{a}$** | **109.8$^{c}$** | **77.4$^{c}$** | **59.8$^{bc}$** | **27.2$^{bcd}$** | **86.9$^{b}$** | **31.6$^{bcd}$** | **14.7$^{abc}$** | **47.3$^{cd}$** |
| **CD (5%)** | **2.6** | **4.7** | **1.9** | **5.8** | **2.9** | **7.0** | **3.2** | **2.4** | **3.8** |

Mean values with different superscript lowercase letters indicate significant difference at the 0.05 level.

**Table6: Estimates of mid-parent ($H_{MP}$) and better-parent heterosis ($H_{BP}$) for the traits of the 55 rice hybrids**

| Cross | Grain yield per plant | |
|---|---|---|
| | $H_{MP}$ | $H_{BP}$ |
| SUR×LAL | 10.64** | 8.94** |
| SUR×BOR | 10.70** | 8.85** |
| SUR×MAY | 6.72** | 5.94** |
| SUR×25B | 13.51** | 11.94** |
| SUR×88B | 10.13** | 7.38** |
| SUR×97B | 3.74* | 1.41 |
| SUR×56B | 24.31** | 22.93** |
| SUR×55B | 1.39 | -1.86 |
| SUR×LAC | 29.05** | 27.11** |
| SUR×LUI | 4.01* | 3.17 |
| LAL×BOR | -6.82** | -10.38** |
| LAL×MAY | 5.71** | 3.22 |
| LAL×25B | -4.02* | -7.30** |
| LAL×88B | 22.20** | 17.74** |
| LAL×97B | 3.99* | 3.37 |
| LAL×56B | 15.01** | 14.68** |
| LAL×55B | 29.34** | 24.38** |
| LAL×LAC | 18.98** | 15.32** |
| LAL×LUI | 14.62** | 12.06** |
| BOR×MAY | 9.71** | 8.64** |
| BOR×25B | 12.01** | 11.73** |
| BOR×88B | 6.50** | 5.59** |
| BOR×97B | 0.08 | -4.10* |
| BOR×56B | -2.88 | -6.11** |
| BOR×55B | 8.53** | 7.14** |
| BOR×LAC | -0.55 | -0.64 |
| BOR×LUI | -0.51 | -1.52 |
| MAY×25B | 4.36* | 3.57 |
| CD (5%) | 3.53 | 4.08 |
| CD (1%) | 4.66 | 5.39 |

**Table 7: Genetic distance based heterotic grouping of the 11 parental genotypes along with mean yield and heterosis estimates**

| Hybrid category | Frequency of crosses | GD | GYP (g Plant$^{-1}$) | $H_{MP}$ (%) | $H_{BP}$ (%) | $H_{SP}$ (%) |
|---|---|---|---|---|---|---|
| **Summarized by hybrid groups** | | | | | | |
| **G1×G1** | 0.02 | 0.29 | 28.48 | 51.74 | 43.56 | 44.51 |
| **G2×G2** | 0.38 | 0.40 | 26.53 | 37.41 | 23.51 | 34.58 |
| **G3×G3** | 0.02 | 0.19 | 33.76 | 46.05 | 36.00 | 71.26 |
| **G1×G2** | 0.25 | 0.55 | 26.59 | 38.40 | 28.39 | 34.90 |
| **G1×G3** | 0.07 | 0.48 | 25.02 | 20.20 | 10.06 | 26.93 |
| **G2×G3** | 0.25 | 0.55 | 34.61 | 63.70 | 46.15 | 75.60 |
| **Summarized by inter-and intra-group hybrids** | | | | | | |
| **Inter-group** | 0.42 | 0.53 | 28.74 | 40.77 | 28.20 | 45.81 |
| **Intra-group** | 0.58 | 0.29 | 29.59 | 45.07 | 34.36 | 50.12 |
| **Summarized by parental groups involved in hybrids** | | | | | | |
| **G1** | 0.34 | 0.44 | 26.70 | 36.78 | 27.34 | 35.45 |
| **G2** | 0.88 | 0.50 | 29.24 | 46.51 | 32.68 | 48.36 |
| **G3** | 0.34 | 0.41 | 31.13 | 43.32 | 30.74 | 57.93 |

**Table 8: Mean performance of the hybrids in GD-based heterotic groups for the different traits**

| Character | G1×G1 | G2×G2 | G3×G3 | Intra-group mean | G1×G2 | G1×G3 | G2×G3 | Inter-group mean |
|---|---|---|---|---|---|---|---|---|
| **Seedling** | 20.05 | 20.24 | 18.70 | 19.66 | 21.84 | 24.31 | 20.88 | 22.34 |
| **Leaf number** | 3.00 | 2.71 | 2.00 | 2.57 | 2.86 | 2.75 | 3.07 | 2.89 |
| **Seedling establishment** | 67.50 | 79.52 | 85.00 | 77.34 | 71.61 | 83.75 | 83.57 | 79.64 |
| **Days to [illegible]** | 80.25 | 75.04 | 74.25 | 76.51 | 76.91 | 75.00 | 78.13 | 76.68 |
| **Days to 50% [illegible]** | 84.50 | 80.54 | 79.25 | 81.43 | 82.75 | 80.81 | 83.63 | 82.40 |
| **Flag leaf area ($cm^2$)** | 35.52 | 27.16 | 40.25 | 34.31 | 27.77 | 23.40 | 26.51 | 25.89 |
| **Days to maturity** | 111.50 | 108.48 | 109.00 | 109.66 | 111.02 | 107.19 | 110.88 | 109.69 |
| **Culm length (cm)** | 66.15 | 55.86 | 56.98 | 59.66 | 83.08 | 81.56 | 79.39 | 81.34 |
| **Productive tillers $Plant^{-1}$** | 8.50 | 14.60 | 13.50 | 12.20 | 16.43 | 12.66 | 17.91 | 15.67 |
| **Average panicle weight** | 5.80 | 3.31 | 3.38 | 4.16 | 2.89 | 3.37 | 3.03 | 3.10 |
| **Panicle length (cm)** | 29.25 | 25.50 | 23.80 | 26.18 | 27.61 | 25.07 | 23.80 | 25.49 |
| **Grains [illegible]** | 102.18 | 95.97 | 165.35 | 121.17 | 97.19 | 146.90 | 142.48 | 128.86 |
| **Spikelet fertility (%)** | 83.51 | 73.62 | 77.38 | 78.17 | 79.13 | 71.22 | 79.85 | 76.73 |
| **Straw yield $Plant^{-1}$** | 65.95 | 66.77 | 67.99 | 66.90 | 65.46 | 58.37 | 50.15 | 57.99 |
| **Grain yield $Plant^{-1}$** | 28.48 | 26.53 | 33.76 | 29.59 | 26.59 | 25.02 | 34.61 | 28.74 |
| **Biological yield $Plant^{-1}$** | 94.43 | 92.59 | 101.75 | 96.26 | 92.05 | 83.38 | 85.40 | 86.94 |
| **Harvest index (%)** | 30.20 | 28.92 | 32.96 | 30.69 | 29.03 | 30.74 | 40.56 | 33.44 |
| **Amylose content (%)** | 19.80 | 15.10 | 11.05 | 15.32 | 14.62 | 14.94 | 14.26 | 14.61 |
| **Gel [illegible]** | 31.25 | 41.07 | 65.00 | 45.77 | 50.93 | 39.38 | 48.07 | 46.13 |

| Variables | GYP (g Plant$^{-1}$) | $H_{MP}$ (g Plant$^{-1}$) | $H_{MP}$ (%) | $H_{BP}$ (g Plant$^{-1}$) | $H_{BP}$ (%) | $H_{SP}$ (%) |
|---|---|---|---|---|---|---|
| $H_{MP}$ (g Plant$^{-1}$) | 0.9713** | | | | | |
| $H_{MP}$ (%) | 0.9277** | 0.9857** | | | | |
| $H_{BP}$ (g Plant$^{-1}$) | 0.9514** | 0.9890** | 0.9789** | | | |
| $H_{BP}$ (%) | 0.9013** | 0.9641** | 0.9834** | 0.9825** | | |
| $H_{SP}$ (%) | 1.0000** | 0.9713** | 0.9277** | 0.9514** | 0.9013** | |
| GD | -0.0067 | 0.0055 | 0.0034 | 0.0351 | 0.0168 | -0.0067 |

**Table 9: Pearson correlation matrix among mean grain yield, combining ability, heterosis and genetic distance estimates for the hybrids**

*, ** Significant at 5% and 1% level, respectively.
